# Short term depression shapes information transmission in a constitutively active GABAergic synapse

**DOI:** 10.1101/370833

**Authors:** Hagar Lavian, Alon Korngreen

**Affiliations:** The Leslie and Susan Gonda Interdisciplinary Brain Research Center, Bar Ilan University, Ramat Gan 5290002, Israel; The Mina and Everard Goodman Faculty of Life Sciences, Bar Ilan University, Ramat Gan 5290002, Israel

**Keywords:** entopeduncular nucleus, globus pallidus, short-term depression, gain control, GABA

## Abstract

Short-term depression is a low pass filter of synaptic information that reduces and flattens synaptic information transfer at high presynaptic firing frequencies. This view questions the relevance of spontaneous high firing rates in some networks. In the indirect pathway of the basal ganglia, spontaneously active neurons in the globus pallidus (GP) form depressing inhibitory synapses in the basal ganglia output nuclei. Using numerical modeling and whole-cell recordings from single entopeduncular nucleus (EP) neurons in rat brain slices we investigated the contribution of the different firing rates of GP neurons to information transmission in the indirect pathway. Wholecell recordings showed significant variability in steady-state depression which decreased as stimulation frequency increased. Modeling predicted that this variability would translate into different postsynaptic noise levels during constitutive presynaptic activity. Our simulations further predicted that individual GP-EP synapses mediate gain control. However, when we consider integration of multiple inputs, the large range of GP firing rates would enable different modes of information transmission in which the magnitude and temporal features of postsynaptic modulation would change as a function of present and past firing rates of GP neurons. Finally, we predict that changes in dopamine levels can shift the action of GP neurons from rate coding to gain modulation. Our results thus demonstrate how short-term synaptic depression shapes information transmission in the basal ganglia in particular and via GABAergic synapses in general.

**Significant statement:** Synapses displaying short-term depression are low-pass filters of synaptic input. Consequently, when presynaptic firing is high, little information passes to the postsynaptic neuron. However, many neurons fire at relatively high frequencies all the time. Are their synapses constantly silenced by depression? We tested this apparent contradiction in the indirect pathway of the basal ganglia, where neurons from the globus pallidus (GP) form constitutively active depressing synapses on basal ganglia output. We hypothesized that the different rates of ongoing activity underlie different modes of action for GP neurons. We show that the synaptic and structural properties of the indirect pathway map GP baseline frequencies to different postsynaptic signals, thus demonstrating how short-term synaptic depression shapes information transmission in the basal ganglia.

## Introduction

Short-term depression plays an important role in neural computation. Synapses that show shortterm depression induce low-pass filtering, adaptation and gain control (Anwar et al. 2017; Chance et al. 1998; Fuhrmann et al. 2002; Chung et al. 2002). Tsodyks and Markram (1997) showed that cortical excitatory synapses displaying short-term depression have a limiting frequency above which changes in presynaptic firing only marginally affect the postsynaptic membrane potential. Information transmission is also tuned by the kinetics of synaptic transmission which affect temporal summation (Banitt et al. 2005). This is more prominent in inhibitory synapses which are characterized by slower kinetics than excitatory synapses. While numerous roles have been suggested for short-term depression and temporal summation in information encoding, the contribution of these processes to the computation of spontaneously active networks is less clear. A study investigating the depressing synapses at the Calyx of Held indicated that the baseline firing rate considerably affects information transmitted as presynaptic activity increases (Hermann et al. 2007). Thus, information transfer via depressing synapses is probably dependent on presynaptic short-term dynamics, presynaptic baseline activity and postsynaptic integration.

The basal ganglia are a group of subcortical nuclei involved in motor, limbic, and cognitive functions (Mink 1996). The entopeduncular nucleus (EP) is one of the basal ganglia output nuclei integrating synaptic information from several pathways within the basal ganglia. During movement, cortical information flows through the basal ganglia via the direct, indirect and hyper-direct pathways, and converges on the EP. In the indirect pathway of the basal ganglia, globus pallidus (GP) neurons form inhibitory depressing synapses in the EP. The neurons in the GP are GABAergic and can be classified into arkypallidal neurons projecting to the striatum, and prototypical neurons projecting to the subthalamic nucleus (STN), and to the basal ganglia output nuclei, the EP and substantia nigra pars reticulata (SNr) (Mallet et al. 2012; Abdi et al. 2015). In the STN, SNr, GP and EP, GABAergic synapses from the GP exhibit short-term depression (Kita 2001; Sims et al. 2008; Connelly et al. 2010; Lavian & Korngreen 2016). The steady-state depression of these synapses increases with the firing rate of the GP neuron (Lavian & Korngreen 2016). In a previous study we obtained paired recordings from GP neurons and showed that despite substantial depression the constitutively active GP-GP inhibitory synapse transmitted information to postsynaptic targets as a result of temporal summation (Bugaysen et al. 2013). Thus, both plasticity and kinetics of synaptic transmission should be considered when investigating information transmission via inhibitory synapses in the basal ganglia.

The GP is classically viewed as a station in the indirect pathway that serves as a source of disinhibition to allow movement suppression. However, this view is challenged by recent findings suggesting that the GP is involved in higher functions such as learning (Schechtman et al. 2016; Gittis et al. 2014). Although more is gradually being learned about the role of the GP in motor and non-motor functions key questions remain unanswered. The first is the fact that prototypical neurons in the GP are spontaneously active and fire over a wide range of frequencies up to 100 Hz (Abdi et al. 2015; Dodson et al. 2015). These neurons change their firing rate during spontaneous movements: approximately half increase their firing rate, and half decrease their firing rate (Dodson et al. 2015). Why do GP neurons fire at such a wide range? How is information transmission during movement affected by the baseline firing rate of GP neurons? Here, we used numerical modeling and whole-cell patch clamp recordings to shed light on the contribution of divergent presynaptic activity on information transmission in the basal ganglia indirect pathway in particular, and via GABAergic synapses in general.

## Materials and methods

### Animals

All procedures were approved and supervised by the Institutional Animal Care and Use Committee and were in accordance with the National Institutes of Health Guide for the Care and Use of Laboratory Animals and the Bar-Ilan University Guidelines for the Use and Care of Laboratory Animals in Research. This study was approved by the Israel National Committee for Experiments in Laboratory Animals at the Ministry of Health.

### *In vitro* slice preparation

Brain slices were obtained from 15-21 day old Wistar rats as previously described (Lavian et al. 2013; Stuart et al. 1993). Rats were killed by rapid decapitation following the guidelines of the Bar-Ilan University animal welfare committee. The brain was quickly removed and placed in ice-cold artificial cerebrospinal fluid (ACSF) containing (in mM): 125 NaCl, 2.5KCl, 15 NaHCO_3_, 1.25 Na_2_HPO_4_, 2 CaCl_2_, 1 MgCl_2_, 25 glucose, and 0.5 Na-ascorbate (pH 7.4 with 95% O_2_/5% CO_2_). In all experiments the ACSF solution contained APV (50 μM) and CNQX (15 μM) to block NMDA and AMPA receptors, respectively (Sigma Aldrich, Cat#A6553 and Cat#C239). Thick sagittal slices (300 μm) were cut at an angle of 17 degrees to the midline on an HM 650 V Slicer (MICROM International GmbH, Germany) and transferred to a submersion-type chamber where they were maintained for the remainder of the day in ACSF at room temperature. Experiments were carried out at 37°C and the recording chamber was constantly perfused with oxygenated ACSF.

### *In vitro* electrophysiology

Individual EP neurons were visualized using infrared differential interference contrast microscopy. Electrophysiological recordings were performed in the whole-cell configuration of the patch clamp technique. Somatic whole-cell recordings were carried out using patch pipettes (4–8 MΩ) pulled from thick-walled borosilicate glass capillaries (2.0 mm outer diameter, 0.5 mm wall thickness, Hilgenberg, Malsfeld, Germany). The standard pipette solution contained (in mM): 140 K-gluconate, 10 NaCl, 10 HEPES, 4 MgATP, 0.05 SPERMIN, 5 l-glutathione, 0.2 EGTA, and 0.4 GTP (Sigma Aldrich, pH 7.2 with KOH). In some experiments the pipette solution was supplemented with 0.2% biocytin (Sigma Aldrich, Cat#B4261) to allow staining of cellular morphology after the experiment. Under these conditions the Nernst equilibrium potential for chloride was calculated to be -69.2 mV. The reference electrode was an Ag-AgCl pellet placed in the bath. Voltage signals were amplified by an Axopatch-200B amplifier (Axon Instruments), filtered at 5 kHz and sampled at 20 kHz. Current signals were filtered at 2 kHz and sampled at 20 kHz. The 10 mV liquid junction potential measured under the ionic conditions here was not corrected for. In voltage-clamp experiments the pipettes were coated with Sylgard (DOW corning).

At the end of the experiments with biocytin in the pipette, slices were fixed in cold 100 mM PBS, pH 7.4, containing 4% paraformaldehyde. After fixation, the slices were washed five times in 0.1% Triton X-100 in PBS for a total of 25 min. Nonspecific binding was blocked with 20% normal goat serum and 0.1% Triton X-100 in PBS for 60 min. Slices were then incubated for 24 hours at 4°C with streptavidin-CY3 (1:1000, Sigma Aldrich, Cat#S6402) diluted in 2% normal goat serum and 0.1% Triton X-100 in PBS. Following incubation, slices were washed five times in 0.1% Triton X-100 in PBS for a total of 25 min. To label nuclei, the slices were incubated with Hoechst 33342 (1:1000, Invitrogen, Cat#H5370) for 10 min and then washed three times in 0.1% Triton X-100 in PBS for a total of 15 min. The slices were then placed on glass slides and dried for 15 minutes before being immersed in a mounting solution (Aqua Poly/Mount, Polyscience Inc., Pennsylvania, USA) and covered with a cover slip. Confocal images were obtained with a Leica SP8 confocal microscope (63x/1.4 N.A. oil objective).

Electrical stimulation was applied via a monopolar 2-3KΩ Narylene-coated stainless-steel microelectrode positioned in the GP. The anode was an Ag–AgCl pellet placed in the bath. Stimulation pulses were biphasic 50-500 μA currents (200 μs cathodal followed by 200 μs anodal phase). To identify the limiting frequency of GP-EP synapses, stimulus trains consisted of 50 pulses delivered to the GP at 1-80 Hz. To analyze changes in membrane potential induced by shifts in the activation frequency, the GP was stimulated with 50 pulses at the conditioning frequency, followed by 10 pulses at the tested frequency, followed by 50 pulses at the conditioning frequency.

### Data analysis

All off-line analyses were carried out using Matlab R2016b (Mathworks) and IgorPro 6.0 (WaveMetrics, RRID:SCR_000325) on a personal computer. Data for each experiment were obtained from at least four rats. All experimental results were pooled and displayed as the mean ± SEM, unless stated otherwise. The peak amplitude of each synaptic response was calculated after subtraction of the membrane potential (in current-clamp experiments) or leak current (in voltage-clamp experiments) preceding the stimulation.

### Simulating short-term synaptic plasticity in ionotropic synapses

The mathematical framework used here to simulate short-term plasticity was taken from (Varela et al. 1997). This model captures physiological results but ignores the biological mechanisms underlying short-term plasticity. It is assumed that the postsynaptic amplitude, *A*, relies on two factors; namely the initial amplitude *A_0_*, and the depression variable *D*.

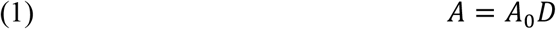

*D* was initially set to 1.

The depression variable *D* was multiplied for each stimulus by a constant *d* representing the depression following a single action potential:

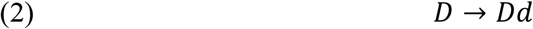

Since *d* ≤ 1, *D* decreased with each action potential. In the model introduced by Varela (1997), after each stimulus, *D* recovered exponentially to 1 using first order kinetics with a time constant *τ_D_*:

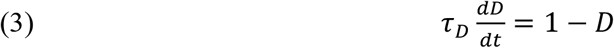

Our previous findings on the GP-EP synapse suggested two time constants for recovery from depression. Thus, the GP-EP synapse was modeled to display short-term depression (*A* = *A_0_ * d*) with *d* = 0.5, *τ_D1_* = 2000 *ms*, *τ_D2_* = 50 *ms*. In simulations of non-plastic synapses, the depression parameter d was set to 1.

### Cellular and Synaptic properties

EP neurons were modeled as:

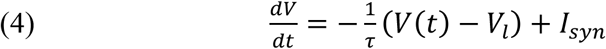

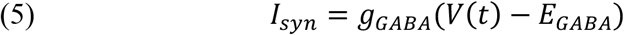

where V is the neuron’s membrane potential, *τ* is the membrane time constant, and I_syn_ is the synaptic current induced by the activity GP neurons. The GABA reversal potential (E_GABA_) was set to -70 mV. The maximal GABA conductance (*g_GABA_*) was set to 10 nS. No voltage-gated channels were implemented in this simulation. Thus, no action potential firing was simulated and all non-linear properties of the membrane of the EP neurons were ignored.

IPSCs were modeled using an alpha function:

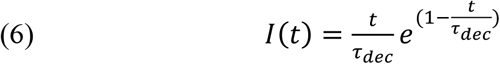

where *τ_dec_* is the time constant of decay of each evoked IPSC, and was set to 10 ms.

### Simulating dopaminergic neuromodulation

Our previous findings showed that dopamine induces a decrease in GP-EP amplitude and a decrease in short-term depression (Lavian et al. 2017). Thus, to simulate an increase in dopamine levels, *g_GABA_* was decreased to 5, and d was increased to 0.7. To simulate a decrease in dopamine levels, *g_GABA_* was set to 15, and d was set to 0.3. These modifications to the model parameters were based on our previous experimental results (Lavian et al. 2017).

## Results

We investigated how short-term dynamics and IPSC kinetics contributed to information transmission in constitutively active GABAergic synapses. Our model system of choice was the synapse between the GP and EP in the indirect pathway of the basal ganglia. We performed voltage-clamp recordings from EP neurons and recorded their response to repetitive stimulation of the GP (Figure 1A-B). As expected, GP-evoked IPSCs displayed short-term depression. Fitting of the evoked currents to an alpha function (eqn. 6) revealed that GP evoked IPSCs were characterized by slow decay kinetics (*τ_dec_* = 9.8 ± 0.2 *ms*, n=20, Figure 1C). It has been shown that the GP-EP synapse displays short-term depression at all frequencies, and that steady-state depression increases exponentially with the presynaptic firing rate (Kita 2001; Lavian & Korngreen 2016). We used these parameters to construct a data driven model of the GP-EP synapse (Figure 1D-F). Figure 1D displays numerically simulated IPSPs evoked by different presynaptic frequencies. The steady-state depression of the GP-EP synapse in our model increased exponentially with the presynaptic firing rate, in a similar manner to that found in slices (Kita 2001; Lavian & Korngreen 2016) (Figure 1E-F).

**Figure 1:**
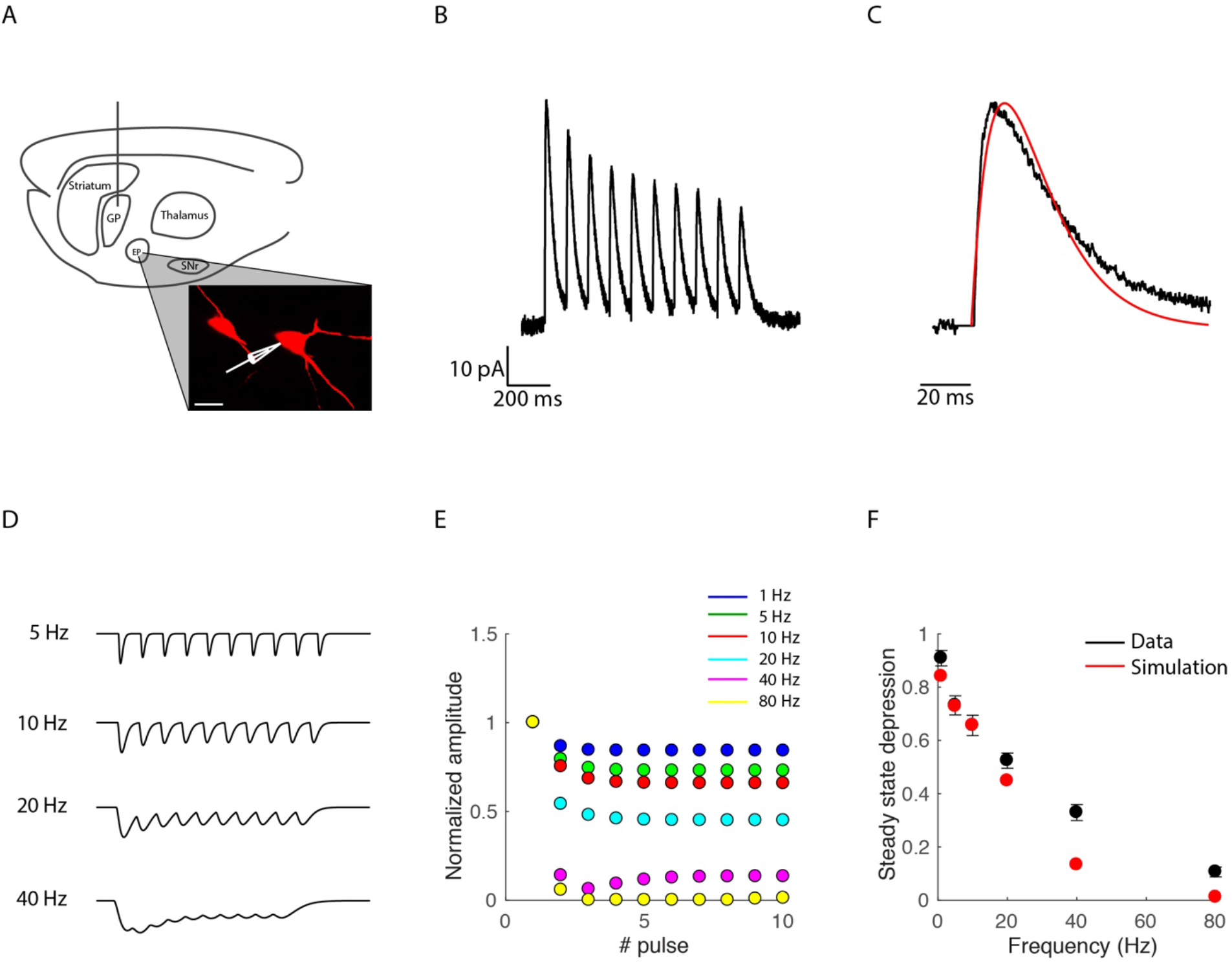
Simulation of GP-EP synapses. A, Experimental design. Illustration of a sagittal brain slice containing the GP and EP. A stimulation electrode was positioned in the GP and whole cell recordings were obtained from EP neurons. A confocal image shows two biocytin filled EP neurons (scale bar = 25 μm). B, Whole-cell voltage-clamp recording displaying the response of an EP neuron to stimulation of the GP at 10 Hz. C, Enlargement of the first evoked IPSC in B. Red curve shows fit of an alpha function (eqn. 6) with a *τ_dec_* of 12 ms. D, simulated traces of GP evoked IPSPs induced by activation of the synapse at 5, 10, 20 and 40 Hz. E, normalized IPSP amplitude evoked by each activation of the simulated GP synapse at different frequencies. F, steady-state depression as a function of the time interval between pulses. Black traces represent population average of the IPSP amplitude evoked by each pulse recorded during GP stimulation at different frequencies. Means ± SEM. Red traces represent data from the simulation.

### Limiting frequency of GP-EP synapses

We used this model to investigate how a GP synapse could change the membrane potential of an EP neuron. Previous studies have shown that depressing synapses can be characterized by a limiting frequency above which increases in the presynaptic firing rate are not reflected in the membrane potential of the postsynaptic neuron due to the increase in steady-state depression (Tsodyks & Markram 1997). The limiting frequency of cortical synapses was found to be ~15-25 Hz (Tsodyks & Markram 1997). Interestingly, the limiting frequency of depressing synapses in the chick auditory system was found to be ~250 Hz (Cook et al. 2003), suggesting that different depressing synapses act over different frequency ranges. Thus, we used our model to determine the limiting frequency of GP-EP synapses. We activated the simulated synapse at various frequencies with 50 pulses, which were enough for the IPSP amplitude to reach a steady state. We measured the change in the simulated membrane potential during the steady state and found that the limiting frequency of the simulated synapse was ~45 Hz (Figure 2A). To validate our findings, we obtained current-clamp recordings from EP neurons and recorded the change in membrane potential induced by electrical stimulation of the GP at different frequencies (Figure 2A). We stimulated the GP with trains of 50 pulses at 5-80 Hz and recorded the response of EP neurons in current-clamp mode (n=28). Our electrophysiological recordings matched the simulated data, showing similar changes in the EP membrane potential (Figure 2A). We calculated the limiting frequency of individual synapses and found that it was 44±5 Hz (Figure 2B-D, n=9). Thus, our recordings corroborated our conclusion that the limiting frequency of GP-EP synapses for information transmission is 45 Hz.

**Figure 2:**
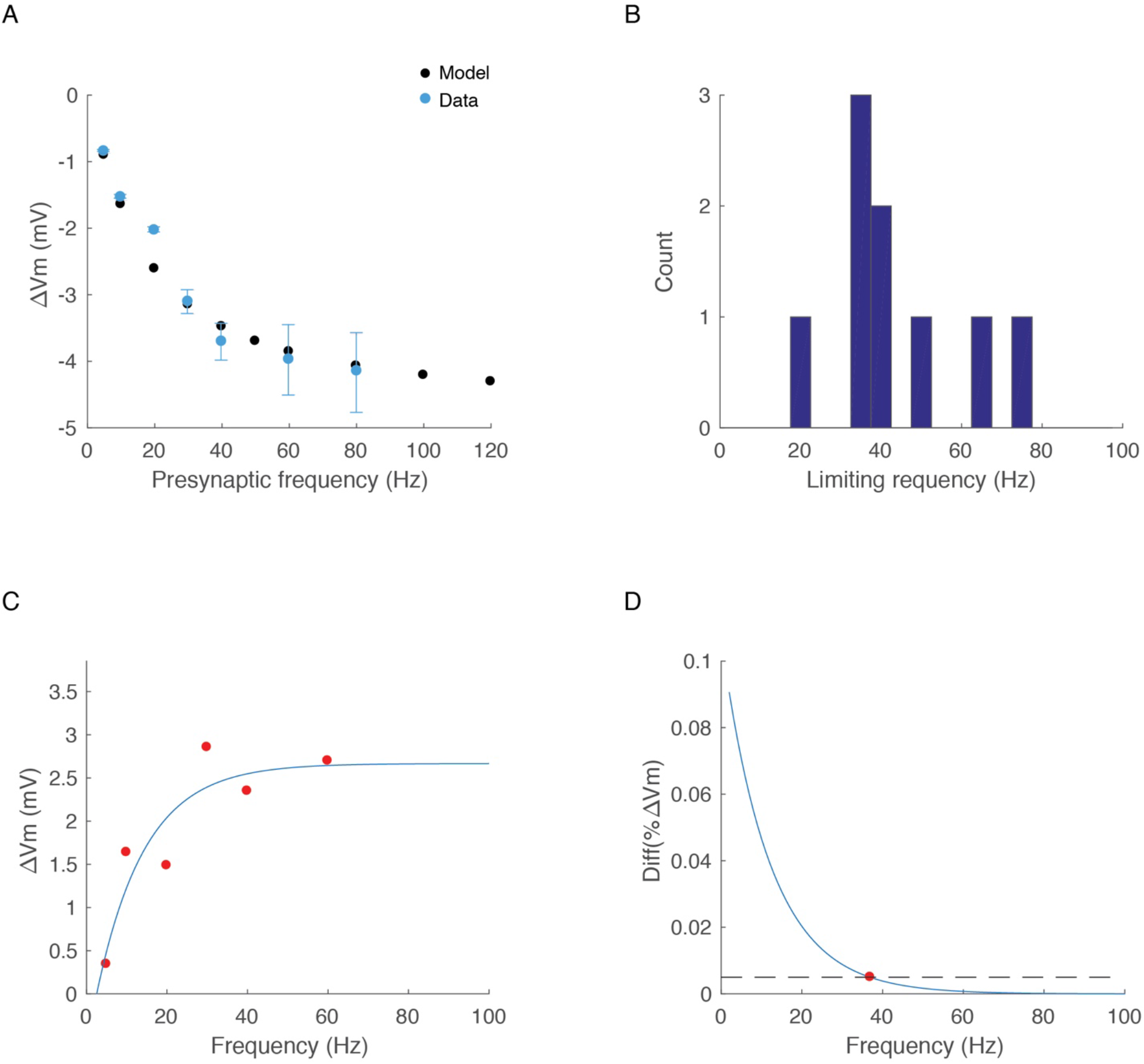
Steady state inhibition induced by GP activity. A, average change in EP membrane potential during steady state inhibition induced by simulated regular stimulation (black) and experimental data (blue). The GP was stimulated at 5 Hz (n=24), 10 Hz (n=24), 20 Hz (n=24), 30 Hz (n=11), 40 Hz (n=9), 60 Hz (n=5) and 80 Hz (n=6). B, distribution of the limiting frequency calculated for 9 individual GP-EP synapses, calculated from GP evoked IPSPs. C-D, example of calculation of the limiting frequency of a GP-EP synapse. C, change of the EP membrane potential as a function of the stimulation frequency of the GP (red dots). The data were fitted with an exponential function (blue trace). D, the limiting frequency was extracted as the activation frequency that induced a change in Vm that was smaller than 5%.

### Baseline firing rate affects the level and variance of GP induced inhibition

Our whole-cell recordings revealed that different GP-EP synapses have different limiting frequencies, with a distribution centered around 45 Hz (Fig. 2B). We examined our data and measured the variance of steady state depression (SSD) for each activation frequency (Figure 3A-B). Our data showed that the variance of SSD decreases as the activation frequency increases. This results from the two main factors affecting SSD: the presynaptic level of depression, which is a function of the depression time constant and the depression factor D, and the postsynaptic temporal filtering, which is a function of the membrane time constant. For a single EP neuron, the properties of GP synapses differ, whereas the membrane time constant does not. Thus, at low frequencies, the effect of temporal summation is small and the SSD is mostly a function of presynaptic factors.

**Figure 3:**
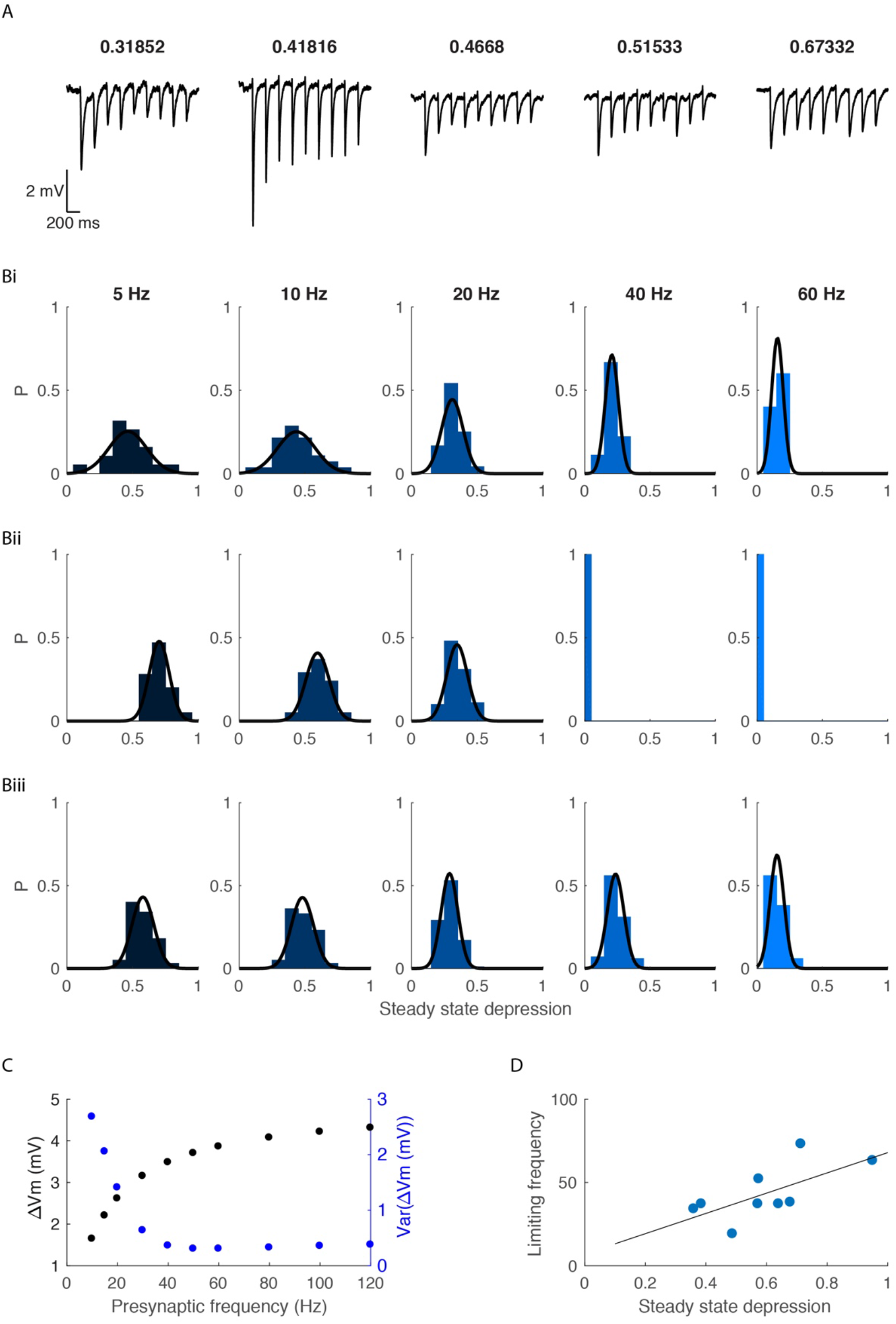
Variability of steady state depression changes as a function of activation frequency. A, example traces from EP neurons during stimulation of the GP at 5 Hz. The steady state depression is indicated above each recording. B, the distribution of the steady state depression for each of the stimulation frequencies extracted from the experimental data (i), from simulated IPSPs evoked by regular stimulation (ii) and from simulated IPSPs evoked by Poisson stimulation (iii). C, the mean and variance of the change in EP membrane potential as a function of the activation frequency. D, the limiting frequency of GP-EP synapses as a function of the steady state depression (measured from stimulation of the GP at 10 Hz). R^2^ = 0.44, p<0.05.

However, as the presynaptic firing increases, the effect of temporal summation on depression increases, and the resulting SSD does not vary. When we added variance to the model, we found that similar to our experimental results, the variance decreased as the activation frequency increased (Figure 3B-C). As expected, the limiting frequency of the GP-EP synapses changed as a function of the steady state depression (Figure 3D). These findings suggest that the baseline firing rate of a GP neuron affects not only the membrane potential of EP neurons, but also the variability of the membrane potential across the EP population.

### Gain control in the indirect basal ganglia pathway

GP neurons are spontaneously active and show brief changes in firing rate during movement (DeLong 1971; Dodson et al. 2015). Thus, a more realistic activity pattern would be one that shifts between frequencies. Previous studies have shown that depressing synapses can mediate gain control such that the change in activity is transmitted with no dependence on the net change in firing rate (Abbott et al. 1997). We used our model to investigate how the baseline activity of a GP neuron could influence information transmission to the EP. We activated the simulated GP synapse at different frequencies, induced a transient change in the activation frequency, and measured the induced change in inhibition (Figure 4A). We quantified the change in inhibition during the transient change by measuring the mean change in postsynaptic membrane potential. Our simulations showed that the mean inhibition during the transient frequency change was nearly independent of the baseline frequency, as opposed to a simulation in which the synapse had no short-term depression (Figure 4B-C). This result is in line with a model of gain control. Small changes were observed when the baseline frequencies were lower than 45 Hz, the limiting frequency of these synapses (Figure Bi, Figure Ci). Thus, acting in isolation, the GP-EP synapse is consistent with a model of gain control where the baseline firing rate of a GP neuron affects postsynaptic inhibition at firing rates that are below the limiting frequency.

**Figure 4:**
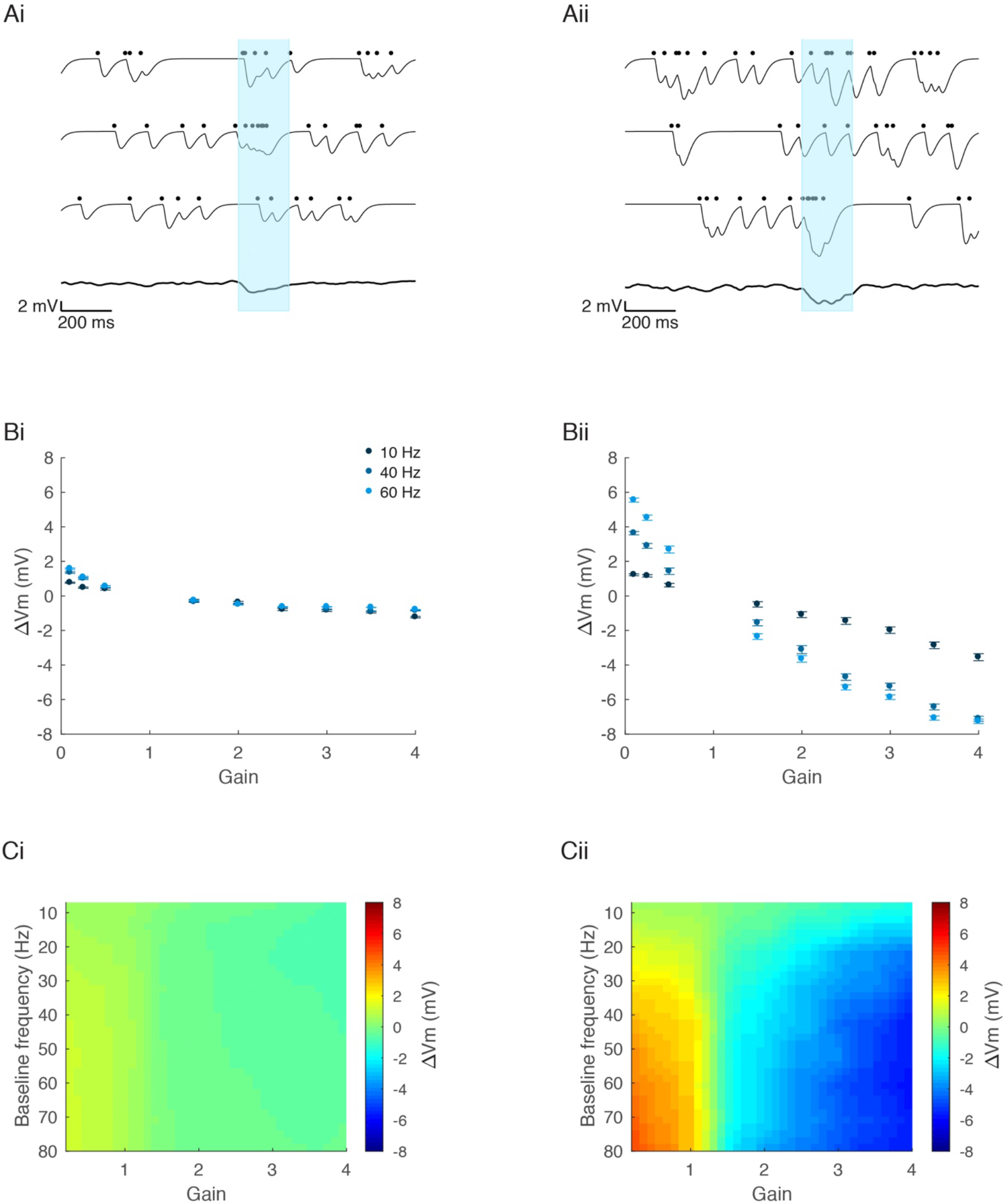
Gain control of the GP-EP synapse. A, simulated traces of GP evoked IPSPs induced by Poisson activation of a depressing synapse (i) and a non-plastic synapse (ii). The synapse was activated at 10 Hz, followed by a transient increase in activation rate to 30 Hz. B, predicted change in membrane potential as a function of the multiplication factor, calculated for baseline frequencies of 10, 40 and 60 Hz. C, predicted change in membrane potential as a function of the multiplication factor and the baseline firing rate.

We carried out whole-cell recordings from EP neurons during electrical stimulation of the GP. The GP was stimulated with 50 pulses at a baseline frequency, followed by 10 pulses at a second activation frequency, and then with additional 50 pulses at the original baseline frequency (Figure 5). Figure 5A shows the change in the membrane potential of an EP neuron during increased activation of the GP. We measured the steady state inhibition during the second frequency and found a very small dependence of the induced inhibition on the baseline frequency, in line with model predictions (Figure 5B-D). Thus, these electrophysiological recording confirm that GP-EP synapses mediate gain control in the indirect pathway.

**Figure 5:**
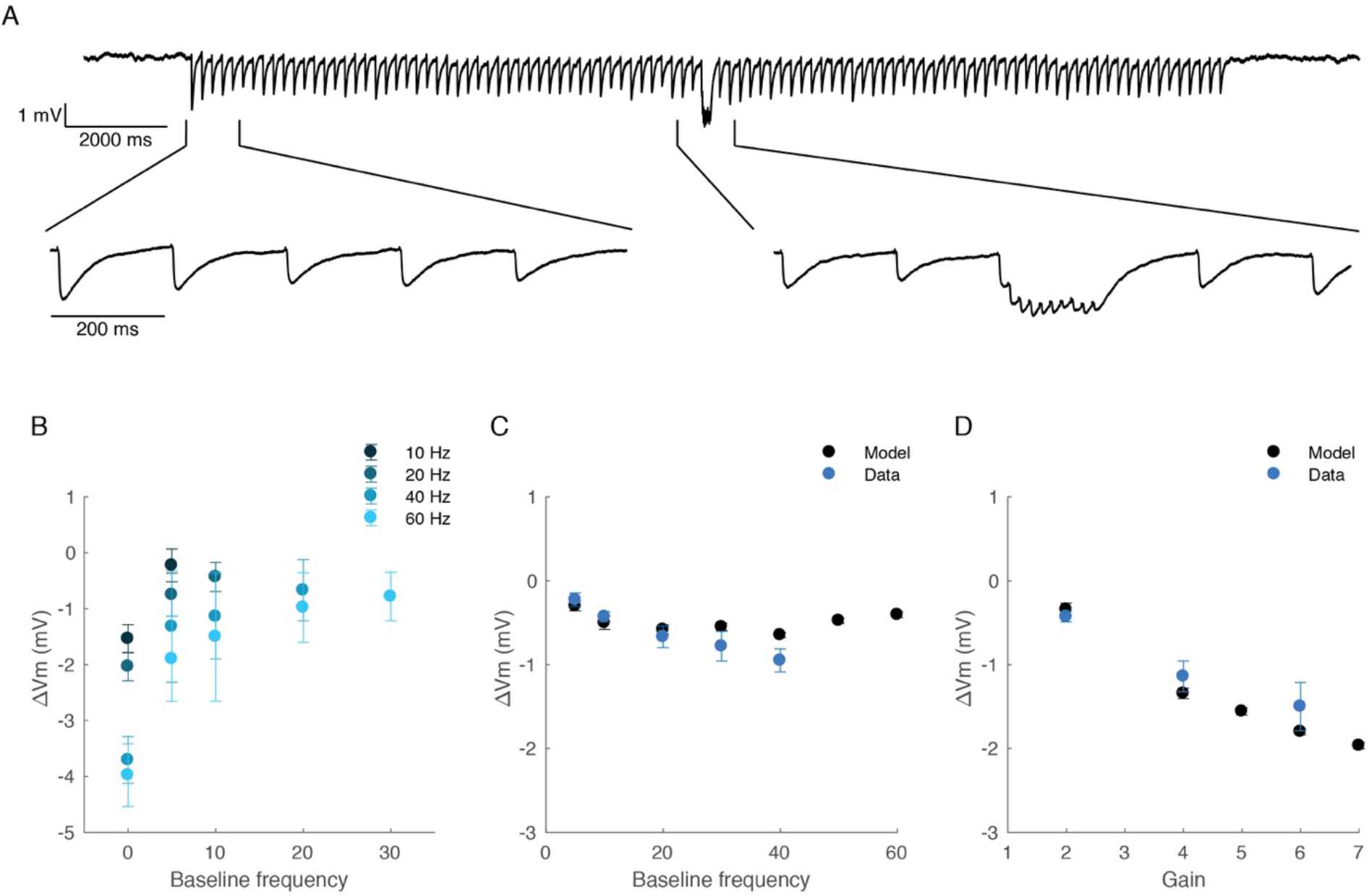
Whole-cell recordings of the response of EP neurons to transient increases in activation frequency of the GP. A, example trace showing the membrane potential of an EP neuron during stimulation of the GP. The GP was stimulated with 50 pulses at 5 Hz, followed by 10 pulses at 60 Hz, followed by 50 pulses at 5 Hz. B, average change in membrane potential as a function of the baseline frequency. The change in membrane potential (ΔVm) was measured during the 2^nd^ activation frequency. C, average change in membrane potential induced by multiplication of the activation frequency by 2 as a function of the baseline frequency. D, average change in membrane potential induced by multiplication of the activation frequency by different gains, baseline frequency of 10 Hz.

### Integration of multiple inputs

Our findings thus far show that the GP-EP synapse appears to mediate gain control. However, there is enormous convergence in the rat indirect pathway, in which 46,000 neurons of the GP converge onto 3200 neurons of the EP, suggesting that each neuron in the EP integrates inputs from multiple GP neurons (Oorschot 1996). In the EP, integration of multiple inputs is expected to be sublinear, since the reversal potential of GABA, which is ~-75 mV in the EP (Lavian & Korngreen 2016), is not too far from the postsynaptic membrane potential. Thus, we used our model to predict how EP neurons would integrate information from multiple GP neurons. Each EP neuron in the model received input from 10 GP neurons that either increased or decreased their firing rate for 200 ms (Figure 6A-B). The EP neurons were also injected with a constant current to ensure a uniform baseline membrane potential of ~-50 mV (Figure 6). We then measured the induced change in postsynaptic inhibition during the second activation frequency (Figure 6C-D). Our simulations demonstrated that during integration of multiple inputs, the induced change in the postsynaptic membrane potential was not only larger, but differentially dependent on the baseline firing rate of the presynaptic neurons (Figure 6B-D). Furthermore, when an EP neuron integrated multiple inputs, the induced change in inhibition depended on the baseline frequency for both high and low gains (Figure 6C-D). Figure 6D depicts the predicted change in membrane potential as a function of both the gain and the baseline frequency.

**Figure 6:**
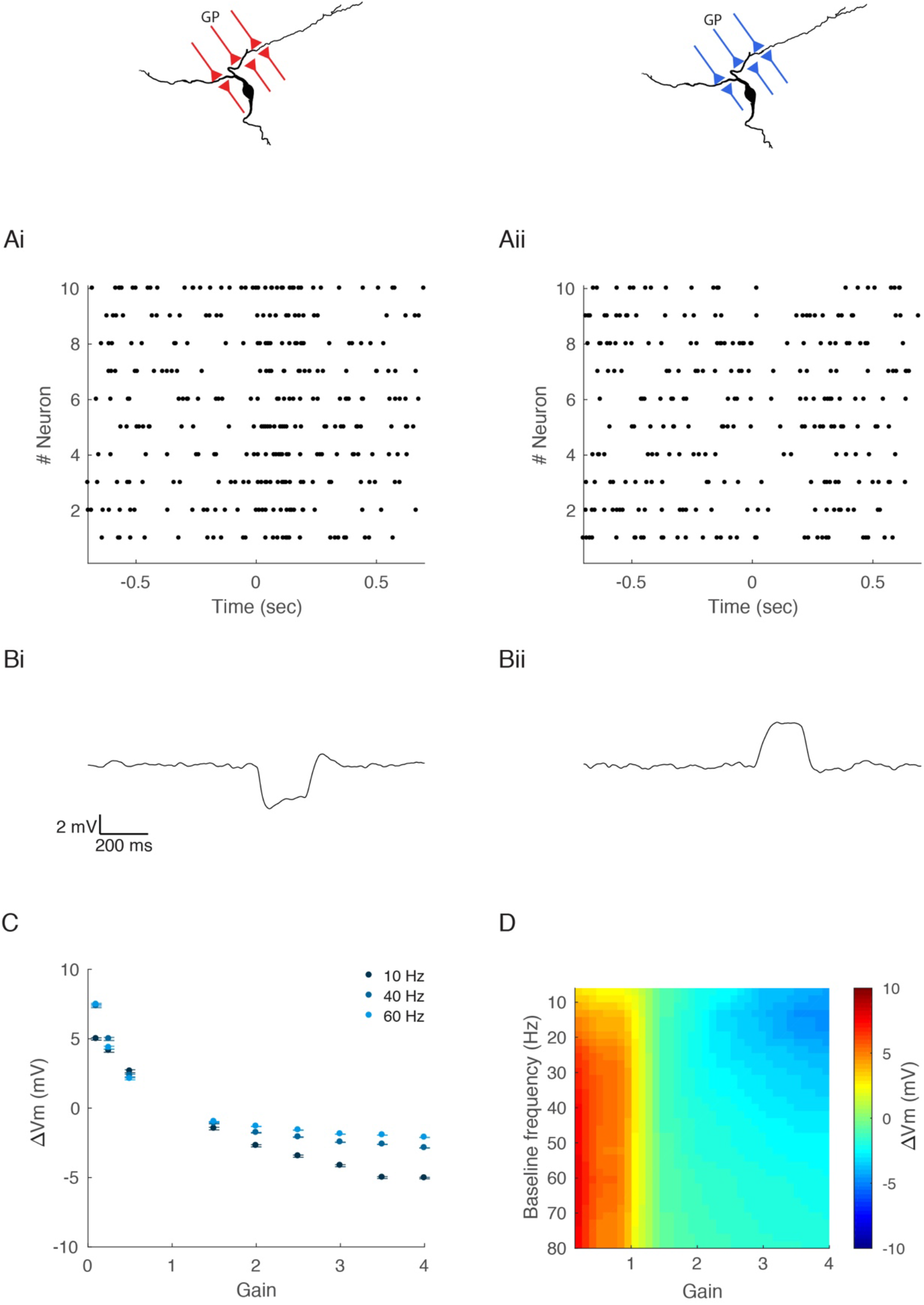
Integration of multiple inputs is mediated by gain control of the GP-EP synapse. GP inputs are labeled red or blue for neurons increasing or decreasing their firing rate, respectively. A, raster plot of the activity of 10 simulated GP neurons that increase (i) or decrease (ii) their firing rate for 200 ms. In these examples the baseline firing rate of all neurons was 20 Hz and the gain was set to 3 (i) or 1/3 (ii). B, predicted membrane potential of an EP neuron that receives input from 10 GP neurons in A. C, predicted change in membrane potential as a function of the multiplication factor, calculated for baseline frequencies of 10, 40 and 60 Hz. D, predicted change in membrane potential as a function of the multiplication factor and the baseline firing rate.

The effect of increased and decreased GP activity on the post-synaptic membrane potential differed in their dynamics. Decreasing GP firing resulted in excitation of the postsynaptic neurons that lasted for the entire duration of change in GP firing. The evoked excitation was constant and did not change significantly during that period (Figure 6Bii). However, increasing the GP firing resulted in a brief inhibition that decayed with time (Figure 6Bi). This result was expected, since GP synapses show short-term depression; hence their inhibitory effect decreases with their activation.

### Integration of multiple heterogeneous inputs leads to several modes of EP modulation

Our findings suggest that GP-EP modulation depends on the number of GP neurons changing their activity and predict different dynamics for EP neurons integrating multiple inputs from GP neurons which either increase or decrease their firing rate. However, EP neurons receive heterogeneous input from the GP, and are innervated by GP neurons that increase their firing rate, as well as GP neurons that decrease their firing rate. Thus, we investigated how EP neurons integrate multiple opposing inputs. To do so, we simulated the change in membrane potential of an EP neuron that received input from 5 GP neurons that increased their firing rate, and 5 GP neurons that decreased their firing rate for an equivalent magnitude (Figure 7A). Our simulations revealed that these opposing inputs did not completely cancel out. Instead, the postsynaptic membrane potential showed a dynamic change depending on the baseline firing rate in the GP (Figure 7B). Integration of heterogeneous inputs resulted in prolonged inhibitory phases when the baseline frequencies were low and prolonged excitatory phases when the baseline frequencies were high (Figure 7C-D). For medium baseline frequencies, integration of heterogeneous inputs resulted in a brief inhibitory phase followed by a brief excitatory phase. The duration and magnitude of the inhibitory phase decreased as the baseline firing rate increased (Figure 7C-D). Conversely, the duration and magnitude of the excitatory phase increased as the baseline firing rate increased. Thus, the combination of several broad changes in GP activity can lead to one of several very different outcomes in the EP. Our simulations predicted a clear mapping from the presynaptic baseline frequency to the type of postsynaptic response. Whereas low and high baseline frequencies resulted in prolonged inhibition or excitation, medium baseline frequencies resulted in temporally focused hyperpolarization followed by depolarization in the membrane potential of an EP neuron. Additionally, the magnitude and temporal component of the GP evoked inhibitory and excitatory phases depended heavily on the baseline firing rate of the presynaptic neurons.

**Figure 7:**
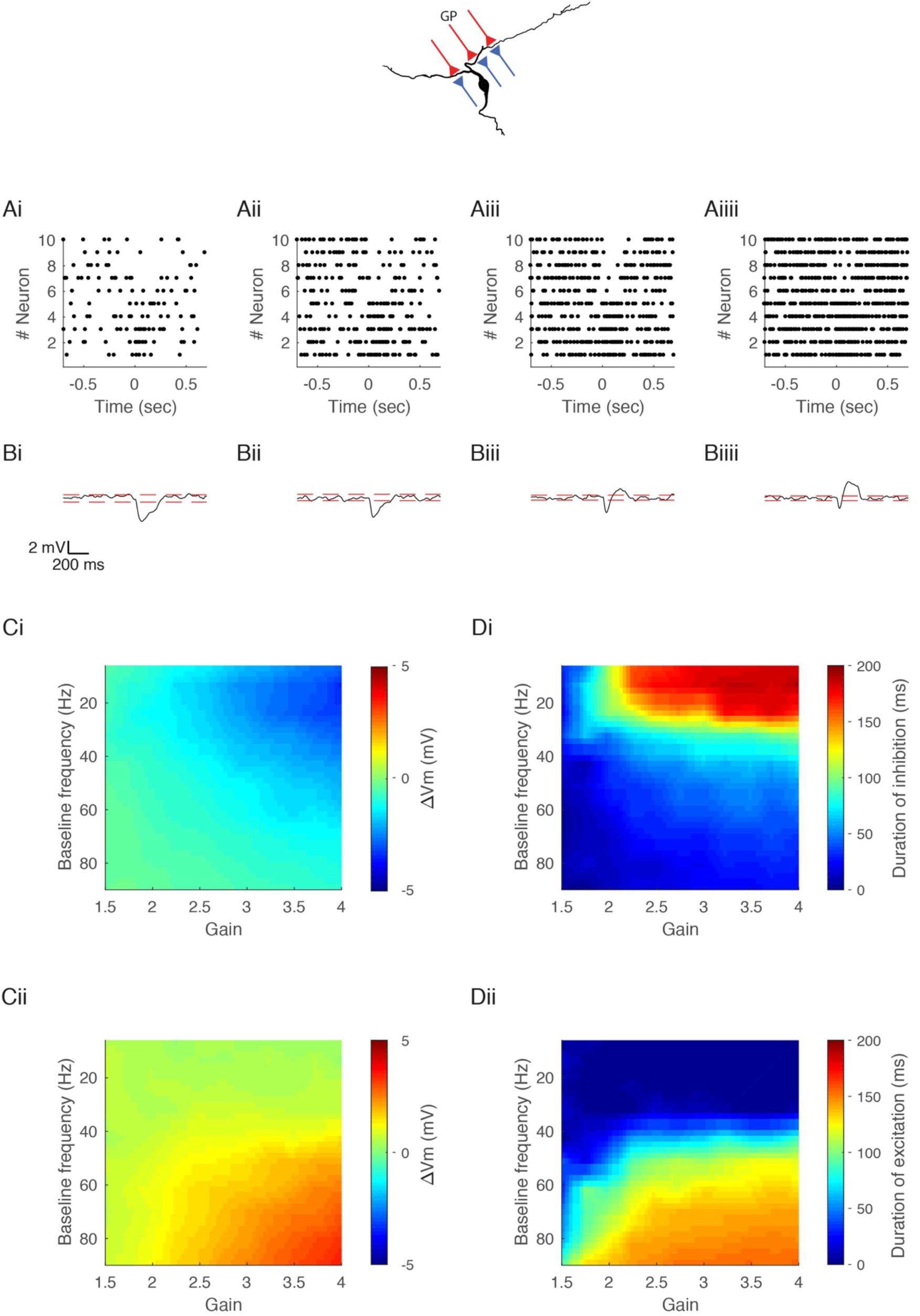
Integration of heterogeneous inputs. GP inputs are labeled red or blue for neurons increasing or decreasing their firing rate, respectively. A, raster plot of the activity of 10 simulated GP neurons that changed their firing rate for 200 ms. Half of the neurons increased their firing rate and half of the neurons decreased their firing rate. The four examples differ in terms of their baseline firing rate: 10 (i), 20 (ii), 40 (iii) and 60 Hz (iiii). B, predicted membrane potential of an EP neuron that receives input from 10 GP neurons in A. Dashed red line indicates two SD below the mean membrane potential. C, predicted change in membrane potential as a function of the multiplication factor and the baseline firing rate. D, duration of the inhibitory phase induced by the change in presynaptic activity. The duration of inhibition was defined as the time during which the postsynaptic membrane potential was significantly hyperpolarized (more negative than 2 SD below the mean membrane potential).

### Dopamine induces a shift in the effects of the GP baseline firing rates

Our findings thus far show that the wide range of baseline firing rates in the GP enables different modes of modulation of neural activity in the EP. These modes of action result from the short-term depression and temporal summation of GP-EP synapses. The shift between the different modes occurs at the limiting frequency of GP-EP synapses. However, the limiting frequency changes as a function of steady state depression, which may change due to neuromodulation. We recently showed that dopamine, a key neuromodulator of basal ganglia function, affects synaptic transmission in the GP-EP synapse by decreasing the amplitude of synaptic transmission and reducing short-term depression (Lavian et al. 2017). Thus, we simulated the effects of changes in dopamine concentration on information transmission from the GP to the EP (Figure 8A-B). Our simulations predicted that changes in dopamine levels would induce a shift in the baseline frequencies that underlie the different modes of action on the EP. An increase in dopamine concentration, simulated by decreased amplitude and reduced depression, should increase the range of baseline frequencies that underlie prolonged inhibition. Conversely, a decrease in dopamine concentration, simulated by an increased amplitude and increased depression, should decrease the range of baseline frequencies that underlie prolonged inhibition (Figure 8C-D). Thus, our simulations predicted that the levels of dopamine would affect the response in the EP and its dependence on GP baseline firing rates.

**Figure 8:**
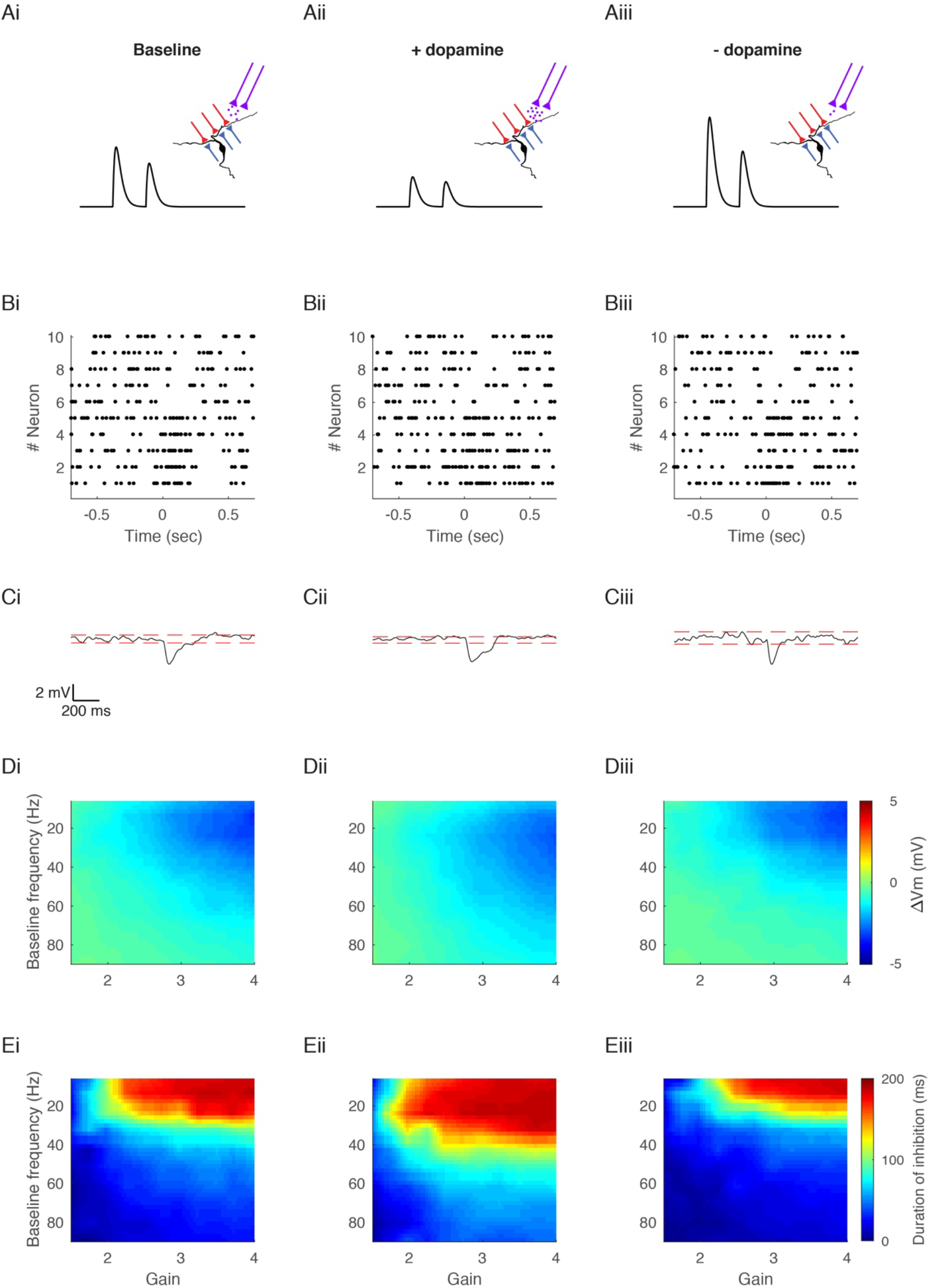
Dopaminergic modulation of integration of pallidal inputs. A, illustration of dopaminergic neuromodulation. GP inputs are labeled red or blue for neurons increasing or decreasing their firing rate. Dopaminergic input from the SNc is labeled in magenta. Black traces illustrate the effect of dopamine on GABAergic conductance of GP-EP synapses. (i), basal dopamine levels, parameters are consistent with the simulations so far. (ii), increased dopamine concentration, simulated by a decrease in the amplitude and depression of GP-EP synapses. (iii), decreased dopamine concentration, simulated by an increase in amplitude and depression of GP-EP synapses. B, raster plot of the activity of 10 simulated GP neurons that change their firing rate for 200 ms. Half of the neurons increased their firing rate and half of the neurons decreased their firing rate. The baseline firing rate of all neurons was 20 Hz. C, predicted membrane potential of an EP neuron that receives input from 10 GP neurons in B. Dashed red line indicates two SD from the mean membrane potential. D, predicted change in membrane potential as a function of the multiplication factor and the baseline firing rate. E, duration of the inhibitory phase induced by the change in presynaptic activity. The duration was defined as the time during which the postsynaptic membrane potential was significantly hyperpolarized (more negative than 2 SD below the mean membrane potential).

### Different onsets of changes in GP activity

Recently, it has been shown that the GP neurons that decrease their firing rate and the GP neurons that increase their firing rate do so at significantly different time scales during spontaneous movements (Dodson et al. 2015). Specifically, neurons increasing their firing rate initiate earlier than those decreasing their firing rate. We thus incorporated a delay of 50 ms in activity into our model (Figure 9A-B). Our simulations predicted that adding a delay would increase the magnitude and duration of both the inhibitory and excitatory phases (Figure 9C-D). Furthermore, incorporating a delay was expected to expand the range of frequencies displaying both phases. For example, with no delay, integration of multiple inputs with a baseline firing rate of 20 Hz can induce a brief inhibition in the EP. However, incorporating a delay of 50 ms induces the emergence of a brief excitatory phase following the inhibitory phase (Figure 9). Thus, incorporating a delay increases the frequency band that induces an inhibition-excitation response in the EP, and reduces the frequency bands which induce prolonged excitation or inhibition.

**Figure 9:**
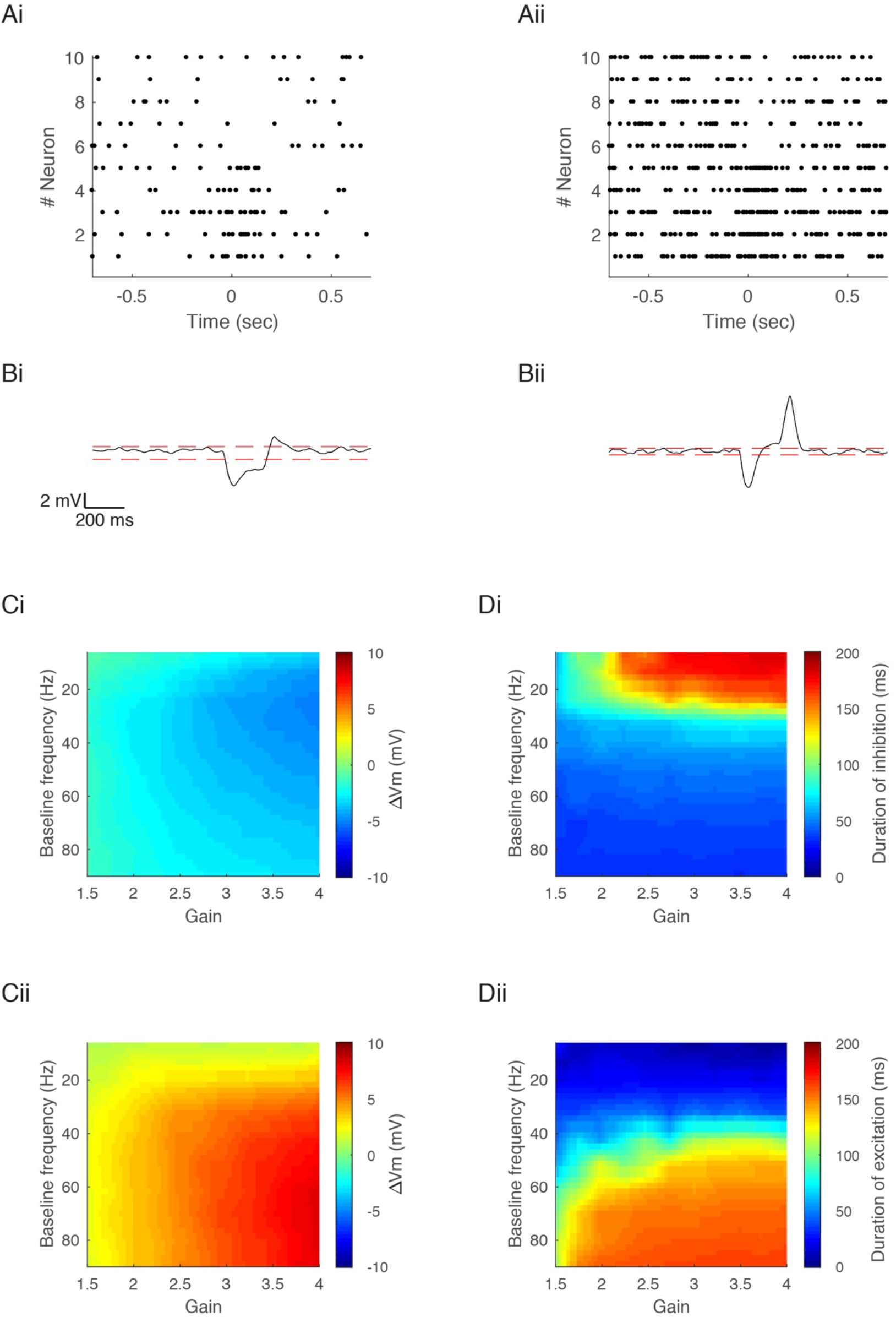
Different delayed firing of increasing or decreasing GP populations evoked short excitation of the EP. A, raster plot of the activity of 10 simulated GP neurons that change their firing rate for 200 ms. Half of the neurons increase their firing rate at t=-50 ms, and half of the neurons decrease their firing rate at t=0 ms. (i), example of low baseline firing rate. All 10 GP neurons have a baseline firing rate of 10 Hz and change their activity by 4/0.25. (ii), example of high baseline firing rate. All 10 GP neurons have a baseline firing rate of 40 Hz and change their activity by 4/0.25. B, predicted membrane potential of an EP neuron that receives input from 10 GP neurons in A. C, predicted change in membrane potential as a function of the multiplication factor and the baseline firing rate during the inhibitory phase (i) or excitatory phase (ii). Calculated as the average change during t=0 ms and t=50 ms (i), and as the average change during t=150 ms and t=200 ms (ii). D, duration of the inhibitory phase (i) and excitatory phase (ii) induced by the change in presynaptic activity. The duration was defined as the time during which the postsynaptic membrane potential was significantly hyperpolarized (i, more negative than 2 SD below the mean membrane potential) or depolarized (ii).

**Figure 10:**
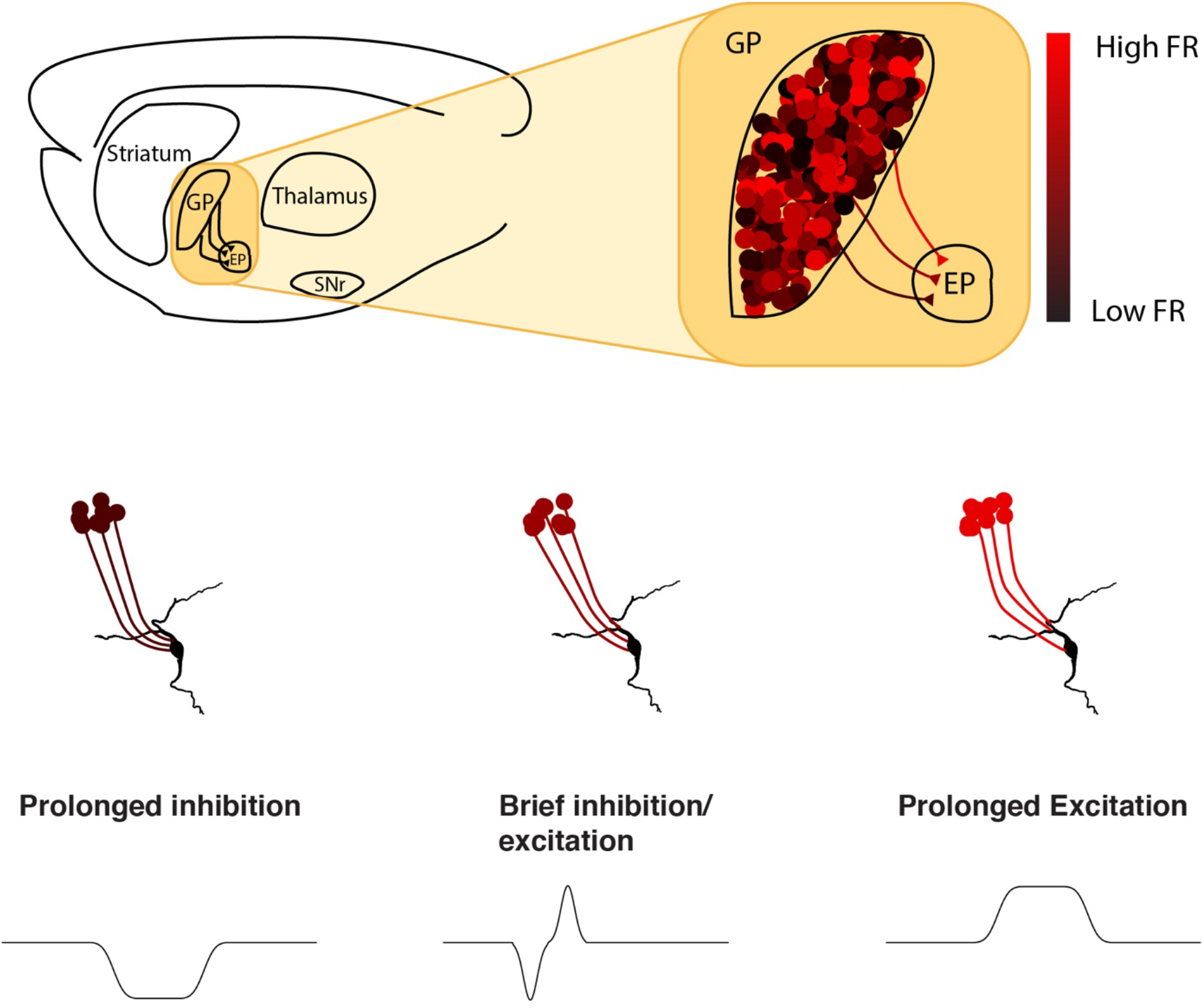
Short-term depression and temporal summation allow different baseline frequencies to underlie different modes of action in the globus pallidus. Upper panel: schematic illustration of the basal ganglia indirect pathway in the rat brain. Neurons in the GP project to the EP and fire at a wide range of frequencies. Lower panel: different modes of action are enabled by the different baseline frequencies. Low baseline firing rate in the GP allows GP neurons to induce prolonged inhibition in the EP, medium baseline frequencies allow GP neurons to induce brief inhibition or excitation in the EP, high baseline frequencies allow GP neurons to induce prolonged excitation in the EP.

## Discussion

We explored the role of short-term depression on information transmission in a constitutively active GABAergic synapse of the basal ganglia’s indirect pathway. We demonstrated that the wide frequency range of GP neurons leads to different levels of inhibition as well as variance in the postsynaptic membrane potential. The average limiting frequency of GP-EP synapses suggests that the inhibitory control of GP over neurons in the EP depends on the baseline firing rate of the GP at frequencies below 45 Hz. During the integration of multiple heterogeneous inputs, different baseline frequencies of GP neurons underpinned different modes of EP modulation. The direction, amplitude and duration of change in postsynaptic membrane potential depended on the baseline firing rate in the GP. Thus, due to short-term depression and slow kinetics of GP-EP synapses, the large range of GP firing rates enabled different modes of information transmission, in which the magnitude and temporal features of postsynaptic modulation changed as a function of present and past firing rates of the neurons in the GP.

The classical view of basal ganglia function suggests that the two main pathways of this system have opposing effects on movement: information flow in the direct pathway facilitates movement, whereas information flow in the indirect pathway inhibits movement. This model has been challenged by numerous findings suggesting that the roles of these two pathways are not as segregated as once presumed. Accumulating evidence shows that a similar amount of direct and indirect striatal spiny projection neurons change their activity in a similar manner during movement (Markowitz et al. 2018; Cui et al. 2013). The GP, classically viewed as a station in the indirect pathway, also receives input from striatal spiny projection neurons of the direct pathway (Parent et al. 1995; Kawaguchi et al. 1990). Moreover, during movement, half of the neurons in the GP increase their activity, and thus do not adhere to their classical role of disinhibition (Dodson et al. 2015). Furthermore, in vivo studies in rodents and primates have shown the involvement of the GP in non-motor functions, such as learning (Gittis et al. 2014; Schechtman et al. 2016). Taken together, the indirect pathway appears to encode multiple types of information, and does not solely serve as a source of disinhibition.

In the indirect pathway of the basal ganglia, GP neurons form GABAergic depressing synapses in the EP. EP neurons are spontaneously active and fire in vivo at ~26 Hz (Ruskin et al. 2002; Benhamou & Cohen 2014). We have previously shown that stimulation of the GP at 40 or 80 Hz can silence spontaneously active EP neurons, and that stimulation of the GP at lower frequencies can induce a transient decrease in firing rate and spike time entropy of the postsynaptic neuron (Lavian & Korngreen 2016; Lavian et al. 2017). Others have shown that single stimulation of the GP in brain slices can synchronize EP neurons (Kita 2001). However, these studies did not consider that GP neurons are spontaneously active, and that GP synapses are consequently substantially depressed. Thus, the ability of GP neurons to synchronize or silence activity of neurons in the EP might be dependent on the baseline activity of the presynaptic neuron. Moreover, these studies did not consider the convergence in this pathway, which is evident from the massive decrease in the number of neurons, as well as from anatomical studies suggesting that each EP neuron receives input from multiple GP neurons (Oorschot 1996; Bevan et al. 1997; Kincaid et al. 1991; Bevan et al. 1998).

### GP neurons introduce continuous noise to the EP

GP neurons fire over a very wide frequency range, but the functionality of this range is not clear (Abdi et al. 2015; Dodson et al. 2015). We have previously shown that, in primates, injection of bicuculline to the GPe leads to an average increase in GPe firing rate from 84 Hz to 113 Hz (Bronfeld et al. 2010). However, in the same animals the average firing rate of GPi neurons did not change, questioning the ability of GPe neurons to inhibit neurons in the GPi (Bronfeld et al. 2010). Thus, the first goal of the current study was to investigate the potential role of the different ongoing firing frequencies in the GP. Our simulations and whole-cell recordings showed that the limiting frequency of GP-EP synapses is ~45 Hz, which suggests that pallidal input can introduce different levels of inhibition to the EP when firing at frequencies below 45 Hz. We further found pronounced variance in steady-state depression, which decreased with the activation frequency of the GP. Thus, the variability between neurons in the EP is likely to decrease as the activity in the GP increases.

### Integration of multiple heterogeneous inputs

During movement, GP neurons briefly change their firing rate. In other systems, synapses displaying short-term depression mediate gain control, such that the change in postsynaptic membrane potential does not depend on the baseline firing rate of the presynaptic neuron (Abbott et al. 1997). In line with gain control model, our simulations and whole-cell recordings show that when a GP neuron changes its firing rate, the induced inhibition manifests very little dependence on the baseline firing rate. However, when several neurons change their activity, the input-output relationship changes. We demonstrated that integration of opposing inputs with equivalent magnitude did not result in zero change but rather led to dynamic change in the postsynaptic membrane potential. The direction, duration and magnitude of the postsynaptic response were shown to change as a function of the gain and the baseline frequency of the presynaptic neuron. Specifically, during integration of heterogeneous inputs, GP neurons firing at low firing rates can induce prolonged inhibition. This postsynaptic inhibition narrows as the presynaptic frequency increases. Over a band of presynaptic baseline frequencies, an excitatory phase appears after the inhibitory phase. As the presynaptic baseline frequency increases, the excitatory phase becomes larger. When the baseline frequency in the GP is high, integration of multiple inputs in the EP induces prolonged excitation. Thus, while individual GP-EP synapses mediate gain control, when we consider the integration of multiple opposing inputs, the synaptic properties lead to a completely different outcome, which allows different baseline frequencies to transmit markedly different signals to the EP.

### Dopaminergic neuromodulation

Dopamine, a key neuromodulator involved in learning, reward and motor control, modulates neural activity throughout the basal ganglia (Beninger 1983; Schultz et al. 1992). Recent findings have highlighted the importance of dopamine at the output level of the basal ganglia. All the dopamine receptors are expressed in the EP on presynaptic and postsynaptic elements (Lavian et al. 2018). In vivo, dopamine gates information flow in the EP, and is necessary for releasing movements (Chiken et al. 2015; Kliem et al. 2007). We have shown elsewhere that dopamine acts in the EP by differentially modulating synaptic transmission from the striatum and GP (Lavian et al. 2017). These findings indicated that blocking dopamine increases basal transmission and depression in the GP-EP synapse (Lavian et al. 2017). Here we suggest that changes in dopamine levels can shift the action of GP neurons from rate coding to gain modulation. In fact, our simulations suggest that changes in dopamine levels result in a shift in the mapping of the baseline presynaptic firing rate to the postsynaptic response. An increased level of dopamine is expected to decrease the level of synaptic depression and thus increase the range of baseline frequencies that can induce prolonged inhibition. Conversely, when the level of dopamine decreases, this frequency band also decreases.

### Coding in the indirect pathway during movement

The EP, an output nucleus of the basal ganglia, forms inhibitory synapses in the thalamus, thus controlling thalamic ability to modulate movement (Carter & Fibiger 1978; Penney & Young 1981). During movement, neurons in the EP integrate inputs from all three basal ganglia pathways. In the direct pathway, EP neurons receive inhibitory input directly from the striatum such that the continuous inhibition from the EP is paused, allowing activation of the thalamus and subsequently the cortex. In the hyperdirect pathway, glutamatergic input from the STN is transmitted directly to the EP, increasing thalamic inhibition to prevent or stop movement. Information transmission in the indirect pathway remains less clear, since it travels through multiple stations. As striatal and subthalamic inputs converge in the GP, half of the neurons in the GP are excited and half are inhibited, which raises the question of whether the activity of these neurons affects activity in the EP. We suggest that the different changes in GP activity are in fact transmitted to the EP. We show that due to synaptic and network properties, integration of either homogeneous or heterogeneous inputs can result in several significant changes in postsynaptic excitability, which vary in direction, duration and magnitude.

## Acknowledgments

This work was supported by an individual grant from the Israel Science Foundation to AK (#168/16). The authors declare no competing financial interests.

## References

Abbott, L.F. et al., 1997. Synaptic depression and cortical gain control. Science, 275(5297), pp.220–224.

Abdi, A. et al., 2015. Prototypic and Arkypallidal Neurons in the Dopamine-Intact External Globus Pallidus., 35(17), pp.6667–6688.

Anwar, H. et al., 2017. Functional roles of short-term synaptic plasticity with an emphasis on inhibition. Current Opinion in Neurobiology, 43, pp.71–78.

Banitt, Y., Martin, K.A. & Segev, I., 2005. Depressed responses of facilitatory synapses. J Neurophysiol, 94(1), pp.865–870.

Benhamou, L. & Cohen, D., 2014. Electrophysiological characterization of entopeduncular nucleus neurons in anesthetized and freely moving rats. Front Syst Neurosci, 8, p.7.

Beninger, R.J., 1983. The role of dopamine in locomotor activity and learning. Brain Res, 287(2), pp.173–196.

Bevan, M.D. et al., 1998. Selective innervation of neostriatal interneurons by a subclass of neuron in the globus pallidus of the rat. J Neurosci, 18(22), pp.9438–9452.

Bevan, M.D., Clarke, N.P. & Bolam, J.P., 1997. Synaptic integration of functionally diverse pallidal information in the entopeduncular nucleus and subthalamic nucleus in the rat. J Neurosci, 17(1), pp.308–324.

Bronfeld, M. et al., 2010. Bicuculline-induced chorea manifests in focal rather than globalized abnormalities in the activation of the external and internal globus pallidus. Journal of neurophysiology, 104(6), pp.3261–75.

Bugaysen, J., Bar-Gad, I. & Korngreen, A., 2013. Continuous modulation of action potential firing by a unitary GABAergic connection in the globus pallidus in vitro. J Neurosci, 33(31), pp.12805–12809.

Carter, D.A. & Fibiger, H.C., 1978. The projections of the entopeduncular nucleus and globus pallidus in rat as demonstrated by autoradiography and horseradish peroxidase histochemistry. J Comp Neurol, 177(1), pp.113–123.

Chance, F.S., Nelson, S.B. & Abbott, L.F., 1998. Synaptic depression and the temporal response characteristics of V1 cells. J Neurosci, 18(12), pp.4785–4799.

Chiken, S. et al., 2015. Dopamine D1 Receptor-Mediated Transmission Maintains Information Flow Through the Cortico-Striato-Entopeduncular Direct Pathway to Release Movements. Cereb Cortex, 25(12), pp.4885–4897.

Chung, S., Li, X. & Nelson, S.B., 2002. Short-term depression at thalamocortical synapses contributes to rapid adaptation of cortical sensory responses in vivo. Neuron, 34(3), pp.437–446.

Connelly, W.M. et al., 2010. Differential short-term plasticity at convergent inhibitory synapses to the substantia nigra pars reticulata. J Neurosci, 30(44), pp.14854–14861.

Cook, D.L. et al., 2003. Synaptic depression in the localization of sound (vol 421, pg 66, 2003). Nature, 423(6936), p.197.

Cui, G. et al., 2013. Concurrent activation of striatal direct and indirect pathways during action initiation. Nature, 494(7436), pp.238–242.

DeLong, M.R., 1971. Activity of pallidal neurons during movement. J Neurophysiol, 34(3), pp.414–427.

Dodson, P.D. et al., 2015. Distinct Developmental Origins Manifest in the Specialized Encoding of Movement by Adult Neurons of the External Globus Pallidus Distinct Developmental Origins Manifest in the Specialized Encoding of Movement by Adult Neurons of the External Globus Pallid. Neuron, 86(2), pp.501–513.

Fuhrmann, G. et al., 2002. Coding of Temporal Information by Activity-Dependent Synapses. Journal of Neurophysiology, 87(1), pp.140–148.

Gittis, A.H. et al., 2014. New Roles for the External Globus Pallidus in Basal Ganglia Circuits and Behavior. Journal of Neuroscience, 34(46), pp.15178–15183.

Hermann, J. et al., 2007. Synaptic transmission at the calyx of Held under in vivo like activity levels. Journal of neurophysiology, 98(2), pp.807–820.

Kawaguchi, Y., Wilson, C.J. & Emson, P.C., 1990. Projection subtypes of rat neostriatal matrix cells revealed by intracellular injection of biocytin. The Journal of neuroscience : the official journal of the Society for Neuroscience, 10(10), pp.3421–38.

Kincaid, A.E. et al., 1991. Evidence for a projection from the globus pallidus to the entopeduncular nucleus in the rat. Neurosci Lett, 128(1), pp.121–125.

Kita, H., 2001. Neostriatal and globus pallidus stimulation induced inhibitory postsynaptic potentials in entopeduncular neurons in rat brain slice preparations. Neuroscience, 105(4), pp.871–879.

Kliem, M.A. et al., 2007. Activation of nigral and pallidal dopamine D1-like receptors modulates basal ganglia outflow in monkeys. J Neurophysiol, 98(3), pp.1489–1500.

Lavian, H. et al., 2018. Dopamine receptors in the rat entopeduncular nucleus. Brain Structure and Function, 223(6), pp.1–12.

Lavian, H. et al., 2017. Dopaminergic Modulation of Synaptic Integration and Firing Patterns in the Rat Entopeduncular Nucleus. The Journal of Neuroscience, 37(30), pp.7177–7187.

Lavian, H., Ben-Porat, H. & Korngreen, A., 2013. High and low frequency stimulation of the subthalamic nucleus induce prolonged changes in subthalamic and globus pallidus neurons. Frontiers in Systems Neuroscience, 7, p.73.

Lavian, H. & Korngreen, A., 2016. Inhibitory short-term plasticity modulates neuronal activity in the rat entopeduncular nucleus in vitro. European Journal of Neuroscience, 43(7), pp.870–884.

Mallet, N. et al., 2012. Dichotomous organization of the external globus pallidus. Neuron, 74(6), pp.1075–1086.

Markowitz, J.E. et al., 2018. The Striatum Organizes 3D Behavior via Moment-to-Moment Action Selection. Cell, pp.1–15.

Mink, J.W., 1996. The basal ganglia: Focused selection and inhibition of competing motor programs. Progress in Neurobiology, 50(4), pp.381–425.

Oorschot, D.E., 1996. Total number of neurons in the neostriatal, pallidal, subthalamic, and substantia nigral nuclei of the rat basal ganglia: a stereological study using the cavalieri and optical disector methods. J Comp Neurol, 366(4), pp.580–599.

Parent, A., Charara, A. & Pinault, D., 1995. Single striatofugal axons arborizing in both pallidal segments and in the substantia nigra in primates. Brain Research, 698(1-2), pp.280–284.

Penney, J.B. & Young, A.B., 1981. GABA as the pallidothalamic neurotransmitter: implications for basal ganglia function. Brain Research, 207(1), pp.195–199.

Ruskin, D.N., Bergstrom, D.A. & Walters, J.R., 2002. Nigrostriatal lesion and dopamine agonists affect firing patterns of rodent entopeduncular nucleus neurons. J Neurophysiol, 88(1), pp.487–496.

Schechtman, E. et al., 2016. Pallidal spiking activity reflects learning dynamics and predicts performance. Proceedings of the National Academy of Sciences, 113(41), pp.E6281–E6289.

Schultz, W. et al., 1992. Neuronal activity in monkey ventral striatum related to the expectation of reward. J Neurosci, 12(12), pp.4595–4610.

Sims, R.E. et al., 2008. Functional characterization of GABAergic pallidopallidal and striatopallidal synapses in the rat globus pallidus in vitro. Eur J Neurosci, 28(12), pp.2401–2408.

Stuart, G.J., Dodt, H.U. & Sakmann, B., 1993. Patch-clamp recordings from the soma and dendrites of neurons in brain slices using infrared video microscopy. Pflugers Arch, 423(5-6), pp.511–518.

Tsodyks, M. & Markram, H., 1997. The neural code between neocortical pyramidal neurons depends. Proc. Nat. Acad. Sci. USA, 94(January), pp.719–723.

Varela, J.A. et al., 1997. A quantitative description of short-term plasticity at excitatory synapses in layer 2/3 of rat primary visual cortex. J Neurosci, 17(20), pp.7926–7940.

